# Seed banks alter metacommunity diversity: the interactive effects of competition, germination, and survival

**DOI:** 10.1101/2021.04.26.441520

**Authors:** Nathan I. Wisnoski, Lauren G. Shoemaker

**Affiliations:** Wyoming Geographic Information Science Center, University of Wyoming, Laramie, WY, 82071, USA; Botany Department, University of Wyoming, Laramie, WY, 82071, USA

**Keywords:** dormancy, metacommunity, dispersal, seed bank, competition

## Abstract

Dispersal and dormancy are two common strategies allowing for species persistence and the maintenance of ecological diversity in variable environments. However, theory and empirical tests of spatial diversity patterns tend to examine either mechanism in isolation. Here, we developed a stochastic, spatially explicit metacommunity model incorporating seed banks with varying germination and survival rates. We found that dormancy and dispersal had interactive, nonlinear effects on the maintenance and distribution of metacommunity diversity, where scale-dependent effects of seed banks were modified by local competitive interactions and dispersal. The interplay between seed germination and survival regulated the benefits of seed banks for diversity. Our study shows that the role of seed banks depends critically on spatial processes, and that classic predictions for how dispersal affects metacommunity diversity can be strongly influenced by dormancy. Together, these results highlight the need to consider both temporal and spatial storage when predicting multi-scale patterns of diversity.

## Introduction

Classic community theory posits that ecological communities are shaped by the interplay of density-independent processes, intraspecific density dependence, and interspecific interactions (Andrewartha & Birch, 1954; Mittelbach, 2012; Vellend, 2016). Density-independent factors, such as environmental fluctuations, influence the sizes and relative abundances of species in the community independently of population density. In contrast, the effects of density-dependent processes vary with population size (both positively or negatively). Density dependence can influence per capita growth rates via intraspecific interactions (e.g., crowding, Allee effects, etc.) or interspecific interactions (e.g., competition, predation, mutualism) that generate non-independence among species in their population dynamics. The maintenance of biodiversity and species coexistence arises from the interplay of these processes (Tilman, 1982; Chase & Leibold, 2003). For example, coexistence occurs if niche differences among species (i.e. differences in their optimal local abiotic environments) are large enough to overcome differences in fitness (i.e. differences in competitive abilities) (Chesson, 2000b; Adler *et al*., 2007). Local environmental fluctuations may further promote coexistence, for example via the temporal storage effect (Chesson, 2000b).

Missing from this historical framework is the role of regional community dynamics in structuring local dynamics (Ricklefs & Schluter, 1993; Cornell & Lawton, 1992; Loreau & Mouquet, 1999; Harrison & Cornell, 2008). The rise and development of metacommunity theory have substantially addressed this interplay between local and regional dynamics, thereby formalizing our understanding of the multi-scale processes that structure communities by incorporating spatial heterogeneity and dispersal between local communities (Leibold *et al*., 2004; Holyoak *et al*., 2005; Logue *et al*., 2011; Leibold & Chase, 2018). In particular, metacommunity dynamics are regulated by density-independent responses to abiotic conditions across the landscape, density-dependent biotic interactions, and dispersal (Thompson *et al*., 2020). The relative importance of local- and regional-scale processes for structuring ecological communities is primarily regulated by rates of dispersal. For example, dispersal limitation may promote regional diversity if strong competitors are unable to colonize all available habitat and competitively displace species from locally structured communities (Leibold & Chase, 2018). Higher rates of dispersal can facilitate coexistence by enabling species to track changing environmental conditions, but excessively high dispersal can homogenize local communities and lead to the loss of regional-scale diversity in the absence of local stabilizing interactions (Mouquet & Loreau, 2003; Thompson *et al*., 2020). Thus, in addition to coexistence mechanisms that operate at the local scale, dispersal across landscapes can help maintain diversity at larger scales through spatial coexistence mechanisms that arise from heterogeneity in both the abiotic and biotic local environments (Chesson, 2000a; Amarasekare, 2003; Shoemaker & Melbourne, 2016).

However, metacommunity theory tends to emphasize how spatial processes affect diversity, minimizing the role of temporal life-history strategies for coexistence. As such, our understanding of species coexistence in temporally variable environments has expanded in parallel alongside metacommunity theory over the last few decades (Abrams, 1984; Chesson & Case, 1986; Chesson, 1994, 2000b; Levine & Rees, 2004; Adler *et al*., 2006; Chesson, 2018). As a temporal analogue of dispersal, dormancy is a key mechanism that can promote species coexistence in temporally variable environments. Dormant individuals can accumulate into a “seed bank” within a community, buffering species’ population sizes through time against harsh environmental conditions (Cohen, 1966; Venable & Lawlor, 1980; Brown & Venable, 1986). Transitions into and out of the seed bank can influence local community dynamics and may promote species coexistence via the temporal storage effect if species respond differently to environmental fluctuations and have a mechanism for buffering against poor environmental conditions (Warner & Chesson, 1985; Pake & Venable, 1996; Angert *et al*., 2009; Gremer & Venable, 2014).

Although spatial and temporal processes interactively shape community dynamics, the joint effects of dormancy and dispersal have rarely been combined, and rather theories for diversity maintenance tend to focus on a single process (Leibold & Norberg, 2004; Holt *et al*., 2005; Wisnoski *et al*., 2019; Holyoak *et al*., 2020). The implications of dormancy for metacommunities extend beyond the local scale and can have regional effects through interactions with dispersal (Cohen & Levin, 1987; Venable & Brown, 1988; Buoro & Carlson, 2014; Wisnoski *et al*., 2019). Understanding the interactions between dispersal and dormancy in a multi-species context has important implications in applied settings, such as restoration ecology or invasive species management (Box 1). Empirical evidence that dormancy may play an important role in metacommunity dynamics is accumulating from plant (Plue & Cousins, 2018), zooplankton (Brendonck *et al*., 2017), and microbial (Wisnoski *et al*., 2020) communities in nature. For example, the dormant resting stages of zooplankton that inhabit ephemeral rock pools allow them to contend with extreme hydrological variability and regulate community dynamics during inundation and drying phases (Brendonck *et al*., 2017). During wet periods, propagule buoyancy can regulate inter-pool dispersal along hydrological vectors (e.g., flooding that connects nearby pools), while during dry periods, exposed egg banks of dormant propagules can be dispersed among pools by wind (Vanschoenwinkel *et al*., 2008). Despite these recent advances, a comprehensive investigation into how dormant seed banks influence metacommunity diversity remains lacking.

### Box 1: Empirical applications of metacommunities with seed banks

Beyond strengthening our theoretical understanding of the processes that maintain biodiversity across spatial scales, integrating seed banks into metacommunity ecology also has wide-ranging empirical applications. Applied ecology has been at the forefront in considering seed bank effects on diversity and community composition. In turn, seed bank theory has contributed to recent advances in biological control (Rees & Hill, 2001; Strydom *et al*., 2017), restoration ecology (Bakker *et al*., 1996; Kiss *et al*., 2018; Ma *et al*., 2019), agriculture (Buhler *et al*., 1997; Menalled *et al*., 2001; Ryan *et al*., 2010), and invasive species management (Gioria & Pyšek, 2016; Strydom *et al*., 2017; Gioria & Pyšek, 2017). Despite the importance of seed germination and survival in applied contexts, theory for the joint effects of dormancy and dispersal on cross-scale diversity patterns is less developed, but presents numerous exciting opportunities for future empirical research.

Research on spatially structured seed banks has uncovered a range of patterns and insights. First, seed banks provide “ecological memory” that moderates the effectiveness of biological control strategies and restoration at the landscape scale. This occurs because germination of viable seeds can reestablish populations, especially when coupled with high dispersal at large spatial scales (Bakker *et al*., 1996). For example, in the Tibetan Plateau, subalpine meadows that had been used for farming for 30 years were left abandoned, allowing up to 20 years of natural regeneration (Ma *et al*., 2019). Even with 30 years of farming, the persistent seed bank remained nearly unchanged, preserving the composition of the pre-disturbance subalpine community. As a result of the long-term persistence of the pre-disturbance community in the seed bank, the aboveground community exhibited high resilience, allowing for the natural recovery of the community to the pre-disturbance state after agriculture was abandoned (Ma *et al*., 2019). However, the seed bank can also preserve a memory of spatial dynamics, such as dispersal limitation or priority effects due to different colonization histories among restoration sites. This may manifest as unexplained variation in restoration success, similar to spatial differences in seed bank dynamics observed in other agricultural systems (e.g., Mahaut *et al*., 2018). The long-term “ecological memory” in persistent seed banks, combined with the capacity for rapid spatial spread via dispersal, suggests that the spatial configuration of aboveground and belowground diversity may be important for promoting successful restorations, either via natural regeneration or through the addition of seed mixtures.

Second, it is common to find differences in diversity or species composition between the seed bank and the aboveground community (Hopfensperger, 2007; Vandvik *et al*., 2016), which suggests the potential for historical contingencies (depending on disturbance history, order of germination, or seed bank composition) that could lead to spatial variation in restoration success or control efficacy. For example, a review of experimental and field studies of grassland seed banks found that, in ecosystems with a disturbance regime shaped by frequent disturbance-recolonization dynamics, such as wetlands, persistent seed banks may be able to promote natural recovery of the aboveground community (Kiss *et al*., 2018). However, ecosystems that lack a frequent history of disturbance, or in communities that contain species with transient seed banks, active measures may be needed for successful restoration, such as direct seed addition (Kiss *et al*., 2018). In restorations that suffer from a lack of diversity, alternative strategies may focus on spatial processes. For example, restored sites may benefit from diversity spillover effects of wind-dispersed species from nearby remnant patches that maintain high diversity (Sperry *et al*., 2019). Sufficiently high rates of spatial dispersal may also be necessary to supplement temporal seed bank dynamics for the maintenance of some specialist species (Plue & Cousins, 2018). Thus, restoration planners should carefully consider the combined effects of spatial dispersal and germination from the seed bank, helping to ensure that restored populations are capable of establishing in intended habitats and tracking favorable environments through both time and space.

Third, efforts to curb the spatial spread of invasive species may also need to combat large seed banks dominated by the invasive. Positive feedbacks that in many cases promote invasiveness could drastically hinder efforts to eliminate invasives. For example, in the South African fynbos biome, a biodiversity hotspot, invasion by several Australian *Acacia* species has threatened the rich native biodiversity and efforts to combat their spread have been costly. *Acacia’s* ability to form large seed banks that facilitate their spread is a major contributing factor to their successful spread (Richardson & Kluge, 2008). Metacommunity models that examine the species traits common to invaders may be crucial for predicting how species spread in a spatial community context and which measures might be effective for controlling their spread. Empirical investigations into the joint spatial and temporal processes that promote or hinder invasive spread may be especially important to reduce the social and economic burdens of invasive species.

Here, we develop a mathematical model to explore the implications of dormant seed banks for metacommunity diversity. In particular, we extend metacommunity theory to examine how and when dormancy helps maintain diversity at local and regional spatial scales. First, we separate dormancy into the processes of seed/propagule germination and seed bank survival to explore whether germination or survival has greater effects on diversity across spatial scales. Second, we examine how dispersal modifies the relative importance of seed germination and survival for the maintenance of diversity. Third, we evaluate how the strength of local competition, and thereby stable versus unstable local coexistence, modifies the effects of dormancy on metacommunity diversity. To evaluate these questions, we develop a spatially-explicit metacommunity model that extends the recent framework of Thompson *et al*. (2020) to include a classic model of seed bank dynamics (Cohen, 1966; Levine & Rees, 2004; Levine & HilleRisLambers, 2009). Our model demonstrates that seed bank dynamics can play an especially important role for the maintenance of regional diversity and modifies classic predictions for the scaling of diversity (e.g., Leibold & Chase, 2018; Thompson *et al*., 2020), depending critically on seed bank survival, germination, and the strength of local coexistence.

## Materials and methods

### Metacommunity model with a seed bank

To address our research questions, we use a discrete time, spatially explicit model of species abundances in a metacommunity with local seed banks (Fig. 1). The total population size of species *i* in patch *x* at time *t* + 1 is given by

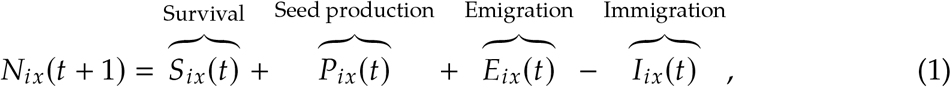

where seed production, *P_ix_*(*t*), is regulated by both density-independent abiotic constraints and density-dependent biotic interactions that determine realized growth, *R_ix_*(*t*), and depend on the germinated fraction of the population, *G_ix_*(*t*). Seed production in a given year and patch are generally modeled as:

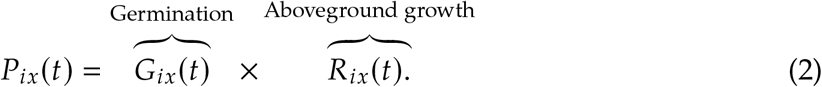

**Figure 1:**
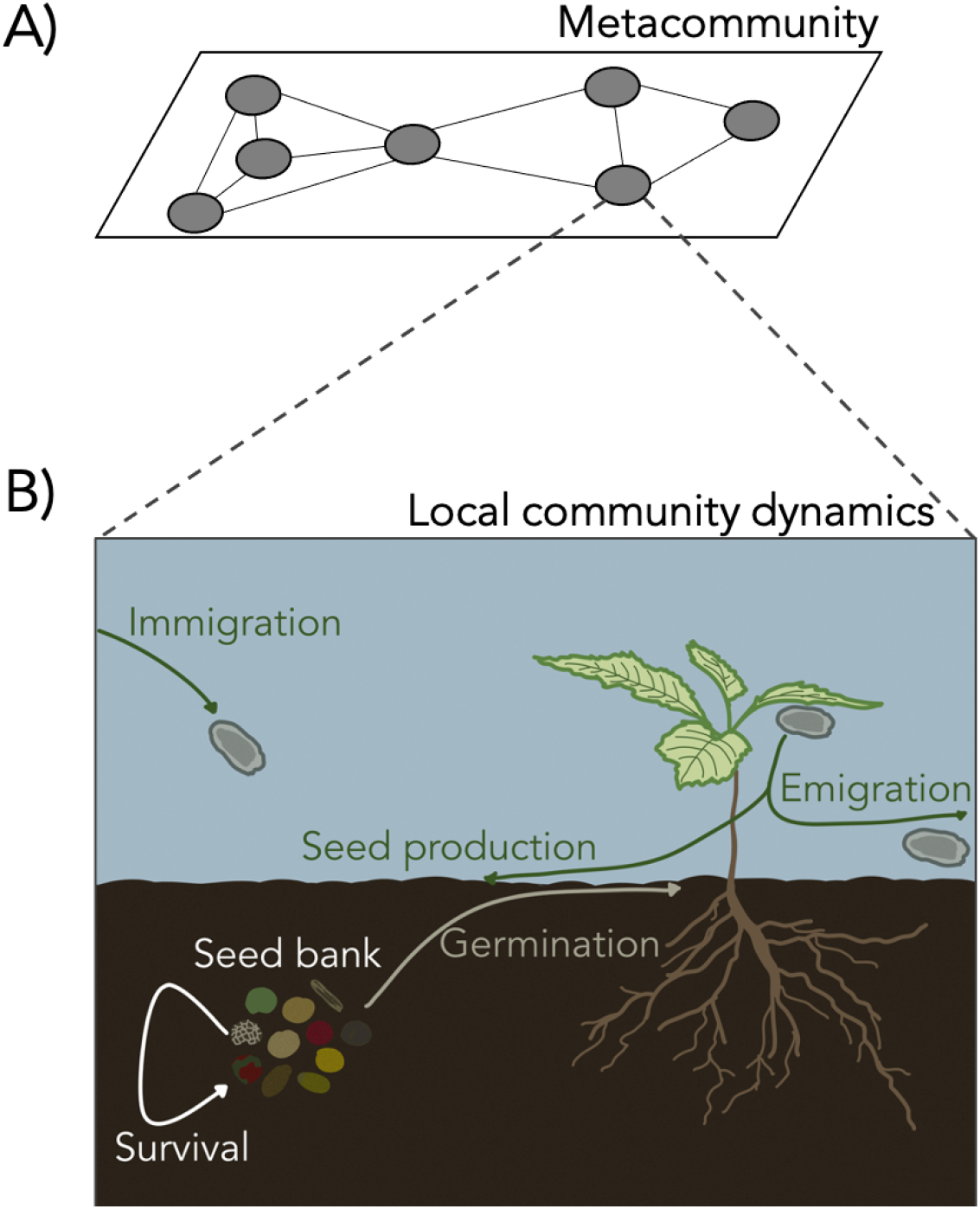
Overview of the metacommunity model. (A) Local communities are uniformly distributed at random across the landscape. In the model, we simulated 100 patches (shown here as gray ovals). For simplicity, lines connecting local communities indicate strong routes of dispersal within the metacommunity (all patches are potentially connected in the model, but nearby patches are more likely to exchange individuals via dispersal). (B) Local community dynamics are governed by aboveground seed production, seed bank survival and seed germination, and immigration and emigration with other patches in the metacommunity, with nearby patches having higher connectivity via dispersal of propagules.

Furthermore, seeds that undergo delayed germination survive in the seed bank, *S_ix_*(*t*); and the seeds generated by the aboveground community exhibit spatially explicit emigration, *E_ix_*(*t*), and immigration, *I_ix_*(*t*).

#### Local seed bank dynamics

At the local scale, we model a community with a seed bank by separating the total seed population into a germinating fraction, *G_ix_*(*t*), and a non-germinating fraction, *N_ix_*(*t*) – *G_ix_*(*t*) (Levine & HilleRisLambers, 2009). To reflect the stochastic nature of germination and survival in natural systems, we model these processes as arising from a binomial distribution. The aboveground, germinating fraction of the community is described as

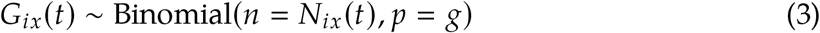

where *N_ix_*(*t*) is the population size of species *i* in patch *x* at time *t*, and *g* is the probability of germination for each individual. The non-germinating fraction, *N_ix_*(*t*) – *G_ix_*(*t*), then survives with probability *s* in the seed bank and is modeled as

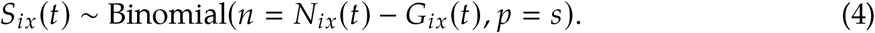

#### Aboveground growth

We determine realized aboveground growth (*R_ix_*(*t*), i.e. per capita production of new seeds) for species *i* in patch *x*, taking into account density-dependent and density-independent limits on population growth of the germinated fraction of the population. We use the classic Beverton-Holt model (Beverton & Holt, 1957) due to its parallel use in both spatial and temporal community ecology theory (Levine & HilleRisLambers, 2009; Shoemaker & Melbourne, 2016; Hallett *et al*., 2019; Thompson *et al*., 2020):

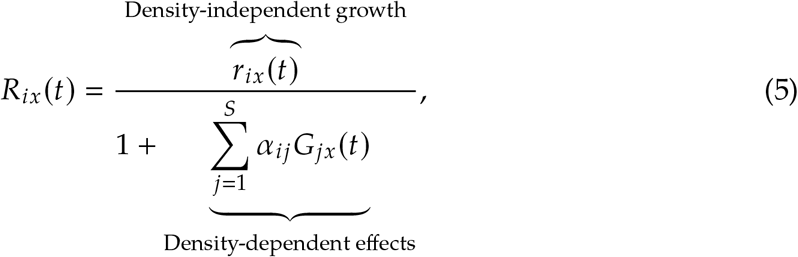

where *α_ij_* is the competition coefficient describing the density-dependent effects of the abundance of species *j* on the growth of species *i*. Note that this summation includes the density-dependent effects of both interspecific (*i* ≠ *j*) and intraspecific (*i* = *j*) competition. We further incorporate density-independent abiotic conditions that affect population growth, *r_ix_*, through a Gaussian function describing species *i*’s niche optimum (*z_i_*) and niche breath (*σ_i_*) in relation to the environmental conditions in patch *x*

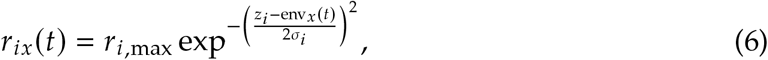

such that species *i*’s maximum growth rate (*r*_*i*,max_) in patch *x* is reduced to *r_ix_*.

To incorporate demographic stochasticity in births to the above equation, we model population size using a Poisson distribution (Poisson (max {*G_ix_*(*t*)*R_ix_*(*t*), 0})), providing integer values for each population or zero if the change in population size leads to local extinction. We incorporate stochasticity throughout our model due to its importance on both population and community dynamics, especially for small population sizes (Lande, 1993; Shoemaker *et al*., 2020).

#### Dispersal

We model the number of emigrants leaving patch *x*, *E_ix_*(*t*), with a binomial distribution

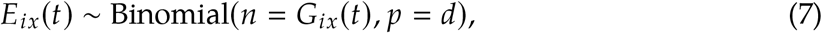

where *d* is the probability of dispersal. Note that dispersal occurs from the germinated portion of the community, *G_ix_*(*t*). The emigrating fraction of species *i* in a metacommunity with *M* patches is given by 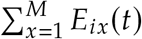. From this pool of emigrants, immigration success in each patch is proportionally determined following a negative exponential dispersal kernel with geographic distance between patches

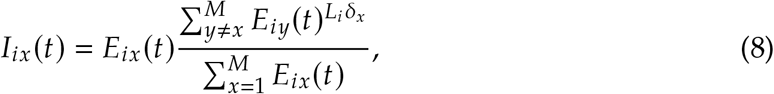

where *L_i_* determines the steepness at which dispersal success decays with geographic distance (*δ_x_*) between patches *x* and *y*.

### Simulations

To investigate (1) the relative importance of germination versus survival on diversity dynamics, (2) how dispersal regulates the effects of germination versus survival, and (3) how the strength of local coexistence and local competition modifies metacommunity dynamics with a seed bank, we ran 30,000 total simulations of our metacommunity model across a wide range of parameter space, as described below.

#### Abiotic conditions

To ensure our results are not contingent upon a given landscape and environmental structure, for each metacommunity simulation, we generated a different landscape structure (i.e., patch connectivity) and environmental conditions. Each metacommunity consisted of 100 patches randomly distributed across a 100 × 100 spatial grid, drawn from a uniform distribution. Spatio-temporal environmental variation was generated anew for each simulation with the “env_generate()” function in the R code provided by Thompson *et al*. (2020) to accompany the revised metacommunity framework that our work extends. To briefly overview, for each patch in the metacommunity, stochastic environmental variables were generated with the RandomFields R package using the “RMexp()” function, and only scenarios with sufficient spatial heterogeneity (i.e., initial environmental differences in the environmental variable greater than 0.6) were kept for simulating metacommunity dynamics. This step ensured that temporal environmental trajectories were spatially autocorrelated, yet sufficiently spatially decoupled across the landscape to support meta-community dynamics.

#### Density-independent abiotic response

To incorporate density-independent responses of different species to environmental conditions, species were assigned niche optima (*z_i_*) evenly distributed in the range [0,1], with equal niche breadth (*σ_i_* = 0.5) among species. Species growth rates under the given environmental conditions in each patch were decreased following the Gaussian function defined above (Eq. 6), such that greater mismatches between species traits and environmental conditions resulted in lower density-independent growth rates.

#### Density-dependence and local coexistence

Density-dependence was incorporated via intra- (*α_ii_*) and interspecific (*α_ij_*) competition coefficients in the Beverton-Holt growth component of the model (Eq. 5). Intraspecific competition was always set to *α_ii_* = 1. We explored two different scenarios to evaluate the implications of locally stable coexistence and competition dynamics versus dispersal and dormancy for diversity dynamics. In *equal intra- and inter-specific competition* (*α_ii_* = *α_ij_*), species coexistence arises from differential responses to abiotic conditions along with dispersal and/or dormancy, as the lack of differences in intra-versus interspecific competition cannot promote coexistence. Alternatively, for *stable competition* (*α_ii_* > *α_ij_*), species can coexist locally in communities due to competitive differences; these processes operate in unison with spatial and temporal coexistence mechanisms arising from dispersal and dormancy. To generate the species interaction matrices, values in the off-diagonal (*α_ij_*) were set to 1 for the “equal intra- and interspecific competition” scenario, and were drawn from a uniform distribution in the range [0,1] for the “stable competition” scenario. The interaction matrix was rescaled by a factor of 0.05 to allow larger population sizes (Thompson *et al*., 2020).

#### Dispersal and dormancy

We simulated our above metacommunity model across a range of parameter values to examine the effect of seed bank germination and survival rates on diversity dynamics. We simulated 10 germination rates, evenly spaced from 10% germination to full germination (i.e., no seed bank) per year (i.e. *g* = [0.1,…, 1]). We also simulated across a range of three survival rates in the seed bank, spanning low (*s* = 0.1), intermediate (*s* = 0.5), and perfect (*s* = 1) survival per year. Last, we simulated across 50 dispersal rates, evenly distributed in logarithmic space (*d* = [10^−5^,…, 1]), ranging from extremely low dispersal (i.e., no metacommunity connectivity; dynamics depend on local processes only) to a well-mixed system with no dispersal limitation between patches (i.e., every individual leaves the patch every year when *d* = 1).

We ran 15,000 simulations each for equal and stabilizing competition coefficients, yielding 10 replicate simulations for each combination of dispersal, germination, and survival rates. We generated a new landscape configuration and new species interaction matrix for each of the 10 replicate simulations.

### Analysis

To quantify changes in aboveground biodiversity across spatial scales, we calculated local (alpha), among-patch (beta), and metacommunity (gamma) diversity for each simulation following a multiplicative partitioning framework:

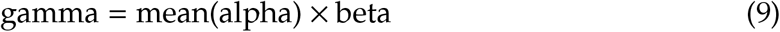

Differences in alpha-, beta-, and gamma-diversity from replicate simulations with the same parameter values illustrate expected variation for a given combination of set dispersal, seed bank survival, and germination rates when considering the combined effects of demographic and environmental stochasticity, landscape configuration, and variation in competition interactions. To assess the overall relationship between dispersal and diversity at different scales, we used local regression (Cleveland, 1979). We visualized trends with locally weighted scatterplot smoothing (LOWESS) computed across all simulations for each parameter set using the ggplot2 R package (Wickham, 2016). Code to reproduce the analysis is available at https://github.com/nwisnoski/metacom-coexistence.

## Results

### Diversity under equal intra- and inter-specific competition

To understand how seed bank dynamics can modify patterns of diversity in the absence of local coexistence mechanisms, we first analyzed a scenario where intra- and interspecific competition were equal. The rate of germination in the seed bank dramatically altered the classic relationship between dispersal rates and alpha-, beta-, and gamma-diversity in metacommunities. With high seed survival, reduced germination shifted the traditional hump-shaped relationship between mean alpha-diversity and dispersal rate, such that dormancy had little effect on alpha-diversity at low dispersal rates (< 10^−3^), but led to strong increases in alpha-diversity with higher dispersal rates (Fig. 2). For example, at high dispersal rates, when the probability of germination was 0.10, local communities had roughly 4 times higher diversity than scenarios without a seed bank.

**Figure 2:**
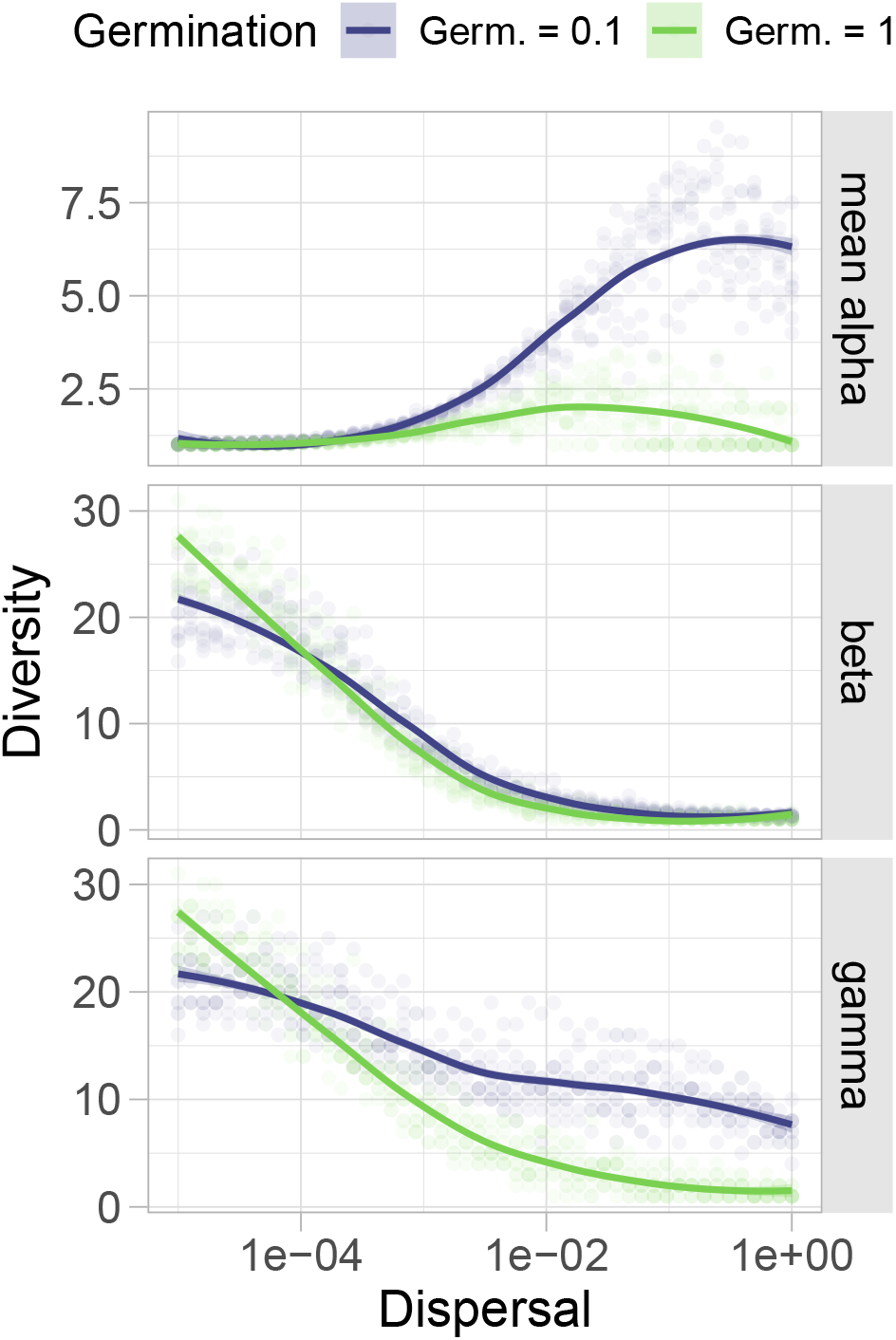
The relationships between mean alpha-, beta-, and gamma-diversity with increasing dispersal are affected by germination rate. Results shown for equal intra- and interspecific competition. Green points and LOWESS fit lines indicate patterns of diversity from traditional metacommunity models (i.e., no dormancy). Blue points and lines indicate patterns that result from the addition of dormancy to metacommunity theory. (Top panel) Reduced germination can shift the expected unimodal relationship between mean alpha-diversity towards the right (so that alpha-diversity peaks at higher dispersal rates) and upwards (so that more diversity overall is maintained within a patch) compared to metacommunities without dormancy. (Middle panel) In this scenario, reduced germination has a smaller overall effect on beta-diversity, but low germination rates can reduce beta-diversity at lower dispersal rates and maintain slightly higher beta-diversity at intermediate dispersal rates. (Bottom panel) Through its effects on alpha- and beta-diversity, reduced germination has important implications for maintaining gamma-diversity. At low dispersal rates, reduced germination leads to losses in gamma-diversity, but once dispersal is sufficiently non-limiting (*d* > 10^−4^) reduced germination can lead to substantially higher gamma-diversity in the metacommunity. For demonstrative purposes, these simulations assumed that survival in the seed bank was high (*s* = 1).

In contrast, reduced germination had a slightly negative effect on beta-diversity when dispersal rates were low (e.g., dispersal < 1 × 10^−4^, Fig. 2). Due to the negative effects on beta-diversity at low dispersal rates and the positive effects on alpha-diversity at intermediate-to-high dispersal rates, persistent seed banks had opposing effects on gamma-diversity at high versus low dispersal rates (Fig. 2). When incorporating seed bank dynamics, as dispersal increases, gamma-diversity no longer declined towards dominance of the metacommunity by a single regionally superior competitor. Instead, seed banks maintained nearly 10 times higher gamma diversity at high dispersal rates. However, at low rates of dispersal (< 10^−4^), reduced germination decreased gamma-diversity relative to scenarios without a seed bank. In this simplified scenario, we focused on large differences in germination rates (0.1 vs. 1), but a fine-grained investigation of germination rate revealed the gradual transitions between these two endpoints (*s* = 1; right column, Fig. 3). Thus far, we assumed full seed bank survival (*s* = 1) to demonstrate the potential effects that reduced germination could have on metacommunity diversity. However, survival rate is likely to be less than perfect.

**Figure 3:**
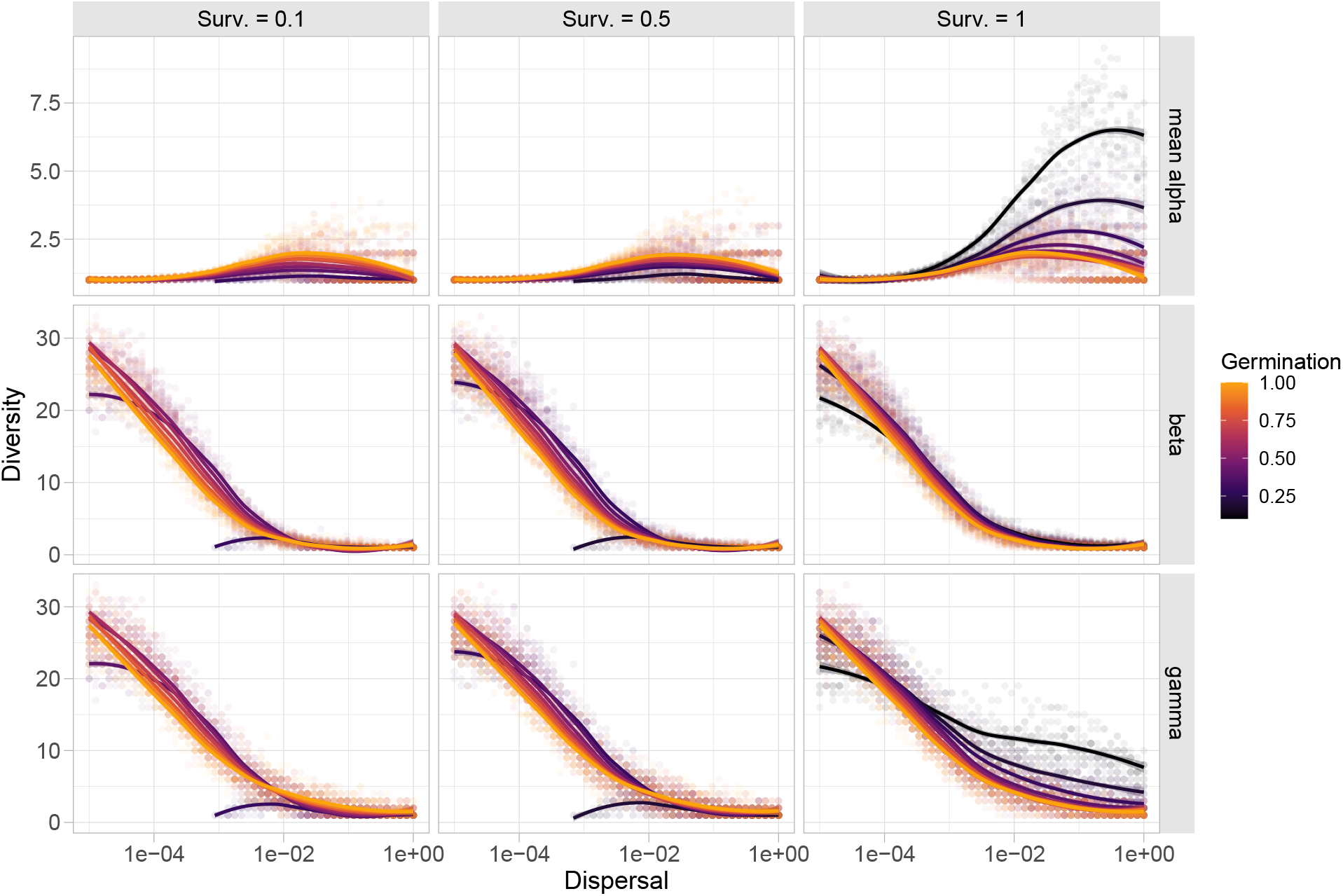
Dispersal-diversity relationships across a range of germination and survival rates with equal competition. In these scenarios, germination plays a key role in shifting the relationships between diversity at different scales and dispersal rate, while survival rate influences the scale-dependent effects of germination and places constraints on the feasible combinations of dispersal and germination that maintain metacommunity diversity. When survival is lower, higher germination rates and higher dispersal rates are necessary to overcome the losses due to reduced survival rates. When survival is high, low germination can reduce gamma-diversity at low dispersal rates, but maintain higher gamma-diversity at higher dispersal rates.

Relaxing our previous assumption and allowing for imperfect survival (*s* < 1), reduced germination was less successful at promoting diversity across the dispersal gradient in the absence of stabilizing competition. Specifically, reduced survival in the seed bank increased the dispersal and/or germination rates necessary for maintaining metacommunity diversity (left and middle columns, Fig. 3). At the lowest germination rates, imperfect seed bank survival also introduced a minimum dispersal threshold (*d* ≈ 10^−3^) necessary for any species to persist; this was most noticeable when germination was less than 0.4 (Fig. S1). Consequently, the highest mean alpha-diversity was detected when dispersal was intermediate and germination was sufficiently high to compensate for the lower survival rates in the seed bank. When germination was higher than the minimum threshold for species persistence, yet lower than complete germination, seed banks maintained higher beta- and gamma-diversity at low dispersal rates. However, because the lowest germination rates were still too low to compensate for the losses associated with reduced survival (darker lines, left and middle columns, Fig. 3), intermediate germination rates maintained regional diversity through positive effects on beta-diversity across much of the dispersal gradient (*d* < 10^−2^). Thus, in metacommunities with low seed bank survival and low germination rates, higher dispersal rates were necessary to allow some populations to persist. However, the lowest germination strategies were no longer as beneficial for the maintenance of diversity, regardless of scale, as they were when seed bank survival was perfect.

### Stabilizing competition coefficients

In natural communities, many species may exhibit niche differences that lead to stabilizing competitive interactions, such as trade-offs in resource requirements. These stabilizing mechanisms can promote species coexistence at local scales, even in the absence of spatial or temporal mechanisms. As such, we extended our analysis above to examine the interplay of dormant seed banks and dispersal on biodiversity with locally stable coexistence via intra- and interspecific competitive differences. When locally stable competitive interactions were included, the effects of germination and seed bank survival strongly differed from patterns without local coexistence mechanisms (Figs. 4, S2). In addition, stabilizing competition yielded differing effects of reduced germination when seed bank survival was high versus low.

**Figure 4:**
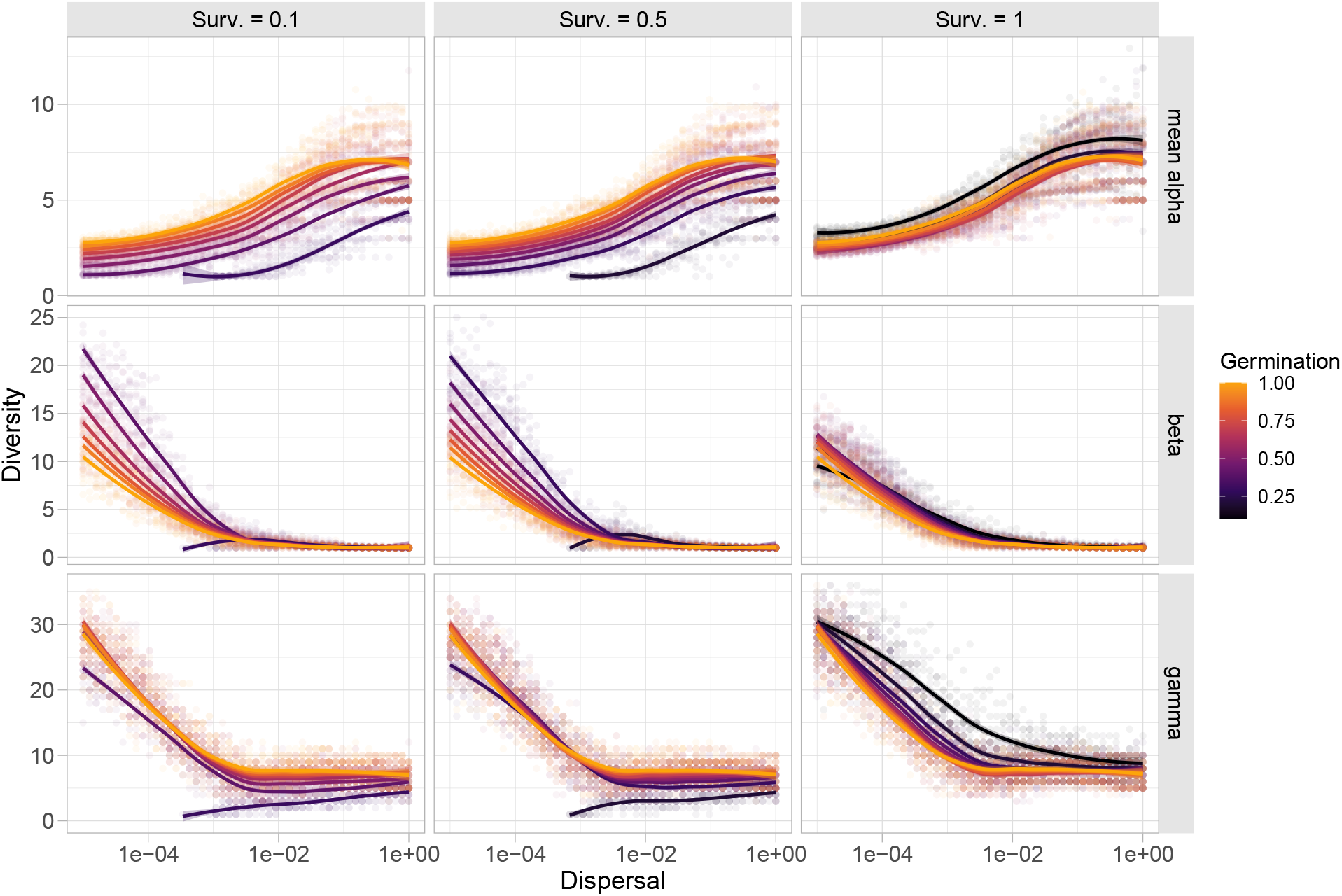
Dispersal-diversity relationships across a range of germination and survival rates with stabilizing competition coefficients. In these scenarios, survival in the seed bank is again a key parameter that regulates the effects of germination at different scales. When survival is low, reduced germination has a negative effect on local diversity by limiting the growth of potentially coexisting species across all dispersal rates, but intermediate germination rates maintain high beta-diversity when dispersal is lower. When survival is high, low germination rates maintain alpha-diversity at all dispersal rates and promote beta-diversity at intermediate dispersal. Consequently, low germination rates maintain high gamma-diversity at all dispersal rates, but especially at low-to-intermediate rates of dispersal.

In the simplifying case where seed bank survival was perfect (*s* = 1, right column of Fig. 4), mean alpha-diversity was an increasing function of dispersal. This positive dispersal-diversity relationship arose because all species could potentially coexist locally due to stronger intraspecific than interspecific competition. Thus, increasing dispersal allowed species to reach all patches where positive growth was possible given abiotic conditions. Interestingly, across all dispersal rates, germination had minimal effects on mean alpha-diversity, except at the lowest germination rates (Fig. 4, S2). When coexistence was locally stable and dispersal was limiting, intermediate germination rates maintained the highest beta-diversity. In contrast, low germination maintained beta-diversity at intermediate dispersal rates. The consequences of reduced germination for diversity maintenance were strongest at the regional scale (Fig. 4, bottom-right panel). In particular, low germination maintained higher gamma-diversity in the metacommunity across the entire dispersal gradient, but the increase in diversity relative to conditions lacking a seed bank were largest at low-to-intermediate dispersal rates.

When seed bank survival was intermediate or low, we observed qualitatively different effects of seed banks on metacommunity diversity (left and middle columns, Fig. 4). With low-to-intermediate seed bank survival, reduced germination had consistently negative effects on mean alpha-diversity across the entire dispersal gradient. The lower the germination rate, the higher the rate of dispersal necessary to maintain diversity at a given survival rate (Fig. S2). In contrast to scenarios with perfect seed bank survival or equal local competition, seed banks had strikingly large positive effects on beta-diversity at low-to-intermediate dispersal and germination rates. Similar to results in the absence of stable coexistence, imperfect survival in the seed bank introduced a minimum threshold for dispersal and germination rates necessary for diversity to persist. When germination rates were above the minimum for persistence, gamma-diversity was less variable across the remaining germination rates. But when dispersal was higher, reduced germination tended to have negative effects on gamma-diversity not through any effects on beta-diversity, but instead by limiting the germination of coexisting species at the local scale when survival was low.

## Discussion

Our results highlight how dormant seed banks affect classic patterns of diversity in meta-communities via interactions among germination, survival, and dispersal. The joint effects of dormancy and dispersal on diversity also depend strongly on whether intra- and interspecific competition are equal or stabilizing (i.e., intra > inter). In the case of equal competition, survival and germination alter the classic dispersal-diversity relationship in several ways (Fig. 3). Lower germination rates increase alpha- and gamma-diversity at higher dispersal rates, but only if survival in the seed bank is sufficiently high; otherwise, reduced germination lowers aboveground alpha- and gamma-diversity. With decreasing seed survival and dispersal, intermediate germination is important for maintaining regional diversity largely through the preservation of beta-diversity at low dispersal rates. With stabilizing local coexistence (Fig. 4), seed bank survival is again an important regulator of the scale-dependent effects of germination on diversity. When seed bank survival is imperfect, any reduction in germination reduces alpha diversity, but intermediate levels of germination preserve beta-diversity if dispersal is not too strong. Yet when seed bank survival is high, reductions in germination increase alpha- and beta-diversity, thereby maintaining higher regional diversity across all dispersal rates. Thus, the nonlinear, scale-dependent effects of dormancy on metacommunity diversity depend on local competitive interactions, as previously highlighted in metacommunity and coexistence literature, but are also strongly dependent on the balance between germination and survival in the seed bank, and the rate of dispersal in the landscape.

### The relationship between seed banks and diversity depends on both local and spatial processes

Much theoretical and empirical research has demonstrated the benefits of seed banks for local diversity maintenance under temporally varying environments (Chesson, 2000b; Saatkamp *et al*., 2014). However, our work indicates that local processes alone may provide an incomplete picture of how seed bank dynamics influence aboveground diversity. In particular, we show that in addition to local scale processes (such as seed bank survival, germination, and the strength of local coexistence), dispersal plays a critical role in regulating the ability of seed banks to maintain locally diverse communities.

Notably, reduced germination and high dispersal can interact to promote alpha-diversity when local stabilizing factors are weak and seed bank survival is high. Previous models lacking dormancy have shown that high rates of dispersal in the absence of local coexistence can reduce diversity by homogenizing the spatial structure of the metacommunity. In other words, high dispersal causes a metacommunity to operate as a single patch favoring superior competitors (Mouquet & Loreau, 2003). Our results indicate that temporal mechanisms associated with seed banks can counteract diversity losses under high rates of dispersal, specifically when competition is equal and seed bank survival is high (Fig. 3). High seed bank survival provides more opportunities for successful germination. The lower the germination rate, the more slowly the stockpile of dormant diversity in the seed bank is depleted (Thompson & Grime, 1979; Thompson, 1987). Consistent with the storage effect, any losses due to poorly timed germination (e.g., during unfavorable environments) are minimized at lower germination rates, but recruitment benefits gained from individuals germinating during favorable environmental conditions replenish the population in the seed bank. Low germination may also reduce aboveground competition and the number of dispersers, further buffering against dispersal-induced diversity loss.

When competitive interactions are stabilizing and seed survival is high, a reduction in germination increases diversity across a broader range of dispersal rates (Fig. 4). Because stabilizing coexistence allows local populations to re-establish from low abundances, even low germination rates are sufficient to promote population persistence following aboveground extinctions. The benefits of reduced germination for local aboveground diversity are consistent across dispersal rates because, even at extremely low dispersal rates where most communities are independent of one another, the seed bank can maintain a stably coexisting community of species favored by the local environment. Hence, at low dispersal rates, higher mean alpha diversity occurred when species could stably coexist (Fig. 4) than when inter- and intraspecific interactions were equal (Fig. 3).

### Local coexistence modifies the dispersal-dependent effects of seed banks on regional diversity

The germination strategies that maximize regional diversity in the metacommunity depend critically on dispersal and the strength of local coexistence. With high survival and no stabilizing competitive interactions, low germination rates can inhibit gamma-diversity when dispersal is limiting and increase gamma-diversity at intermediate-to-high dispersal rates (Fig. 3). In plant communities, for example, dispersal is frequently limiting across a range of ecosystems, even more so in disturbed ecosystems (Myers & Harms, 2009). In metacommunities with local disturbances, previous models suggest that seed banks maintain gamma-diversity at low dispersal rates by preserving both alpha- and beta-diversity (Wisnoski *et al*., 2019). When communities are isolated, our results suggest higher germination rates may be required for some species to persist in local communities, and contribute to beta-diversity, in the face of stochastic population dynamics. Thus, intermediate germination strategies maintain the highest regional diversity at low dispersal rates, but once dispersal is high enough to facilitate environmental tracking across space and time, further reductions in germination maintain significantly higher gamma-diversity.

In contrast, when seeds have high survival rates and competitive interactions are stabilizing, low germination rates promote regional diversity across all dispersal rates (Fig. 4). In this scenario, intermediate germination rates still preserve the highest beta-diversity when dispersal is low. However, because the benefits of reduced germination for alpha-diversity are independent of dispersal, gamma-diversity consistently benefits from the lowest germination rates. Thus, in the case of stabilizing coexistence, at low dispersal rates (< 10^−4^), seed banks contribute to the maintenance of gamma-diversity primarily through their ability to maintain higher mean alpha-diversity in dispersal-limited communities (Fig. 4). However, above this same dispersal rate, both alpha- and beta-diversity were highest when germination is lowest. For example, in fragmented grassland communities across Sweden, the loss of spatial connectivity led to a decline in species reliant on dispersal and impeded the ability of dormancy to maintain some species, but species with long-lived seeds persisted via local seed bank dynamics (Plue & Cousins, 2018). Therefore, the joint benefits of reduced germination for maintaining locally diverse but regionally different communities combine to maintain much higher gamma-diversity across a wide range of intermediate-to-high dispersal values.

### Intermediate germination rates promote diversity under imperfect seed bank survival

When seed bank survival was high, the lowest germination rates often maintained the highest local and regional diversity because remaining in the seed bank was not risky. However, in natural systems, individuals are gradually lost from seed banks due to burial (Bonis & Lepart, 1994; Brendonck & De Meester, 2003), damage (Long *et al*., 2015), and consumption (Janzen, 1971; Horst & Venable, 2018). When seed bank survival is less than perfect, our model highlights how germination rates must be high enough so that individuals germinate before being lost from the seed bank. Otherwise, the seed bank becomes a reproductive sink for aboveground populations. For example, a stochastic population model for the invasive musk thistle *Carduus nutans*, which forms abundant and persistent seed banks with survival surpassing 20 years, showed that the evolutionary stable strategy for germination probability increased with the probability of seed death (Rees *et al*., 2006). That is, germination should be higher when seed survival is low because of the fitness costs associated with losses in the seed bank.

When remaining in the seed bank is risky (i.e., survival = 0.1, 0.5), our model suggests reduced germination decreases alpha-diversity. Natural populations may have insufficient germination rates for many reasons, including recent environmental changes or physiological limitations that prevent optimal germination strategies (Wisnoski *et al*., 2019). For example, in a long-term study of forb communities at the Cedar Creek Natural History Area, experimental nitrogen fertilization caused a compositional divergence between the seed bank and the aboveground community; this discrepancy was hypothesized to arise from germination inhibition (Kitajima & Tilman, 1996). The benefits of intermediate germination for diversity emerged at the regional scale. Previous models suggest that dormancy may be able to substitute for dispersal under certain conditions (Venable & Brown, 1988; Cohen & Levin, 1991; Snyder, 2006), such as when dispersal is limiting and local environments vary through time (Wisnoski *et al*., 2019). Our results emphasize the importance of “temporal dispersal” that maintains beta-diversity when dispersal is low-to-intermediate (< 10^−2^). In contrast, when dispersal is high, low germination and seed survival promote losses from the seed bank, which may explain why, in some cases, gamma-diversity is lower with a seed bank than without one. This loss of regional diversity occurs from the negative interaction of both spatial homogenization and “temporal dispersal” limitation. In the extreme case, with low survival, low germination, and low dispersal, the metacommunity cannot persist.

### Future Directions

Our aim in this study was to develop an understanding of how variation in seed bank dynamics through germination and survival interact with local scale processes (e.g., competition) and regional processes (e.g., dispersal) to affect patterns of diversity. Historically, our understanding of how dormant seed banks can influence patterns of diversity in ecological communities have primarily been informed by local scale studies. Likewise, metacommunity research has largely overlooked the potential role of seed banks in influencing the structure and dynamics of local communities as well as the potential interactions that emerge between dispersal and dormancy that can affect regional biodiversity. Our model demonstrates a range of intuitive yet novel predictions regarding the implications of dormancy in metacommunity theory as well as the role of spatial processes in affecting local seed bank dynamics. Although we have modeled germination, survival, competition, and dispersal as independent traits, covariance among these traits could lead to trait syndromes that have implications for metacommunity dynamics and the maintenance of diversity and present an exciting next direction (Buoro & Carlson, 2014; Rubio de Casas *et al*., 2015; Wisnoski *et al*., 2019). Likewise, we follow similar assumptions from other metacommunity studies by assuming that species exhibit similar dispersal probabilities (Shoemaker & Melbourne, 2016; Thompson *et al*., 2020). Future work investigating trade-offs among dispersal and competition, germination, or survival may reveal favorable strategies that allow species to coexist in a spatio-temporally variable landscape.

## Conclusions

Seed bank dormancy has played a key role in empirical studies of diversity and community turnover, including in restoration settings (Box 1; Saatkamp *et al*., 2014). Simultaneously, it is classically invoked as a key mechanism that promotes coexistence through the temporal storage effect (Warner & Chesson, 1985; Adler *et al*., 2006; Angert *et al*., 2009). Yet, despite this history, its incorporation into metacommunity models has lagged, making it difficult to predict how dispersal and dormancy will alter diversity at local and regional scales. Here, we demonstrate that seed survival and germination interact with dispersal to affect diversity across spatial scales. For example, the combination of high dispersal and low germination can overcome the classic hump-shaped relationship between dispersal and alpha-diversity predicted in many metacommunity models, but only when seed bank survival is high and competitive interactions are neutral. When seed bank survival is low, the seed bank becomes a demographic sink that reduces alpha-diversity. Thus, the implications of dormant seed banks scale nonlinearly with space to influence regional patterns of biodiversity. Integrating insights from both empirical and theoretical studies is likely to be a key step towards understanding the spatial scales at which dormant seed banks promote or erode diversity in natural systems. In particular, empirical estimates of survival rates in the seed bank will be especially informative given that theory predicts survival to be an important regulator of the scale-dependent patterns of biodiversity in metacommunities.

## Acknowledgements

We acknowledge computational support from the Teton Computing Environment (https://doi.org/10.15786/M2FY47) at the Advanced Research Computing Center (ARCC) at the University of Wyoming. This research was supported by the Microbial Ecology Collaborative with funding from NSF award #EPS-1655726.

