## Supplemental Figures for "Seed banks alter metacommunity diversity: the interactive effects of competition, germination, and survival"

April 26, 2021

Nathan I. Wisnoski

Wyoming Geographic Information Science Center
University of Wyoming, Laramie, WY, 82071, USA

Lauren G. Shoemaker

Botany Department
University of Wyoming, Laramie, WY, 82071, USA

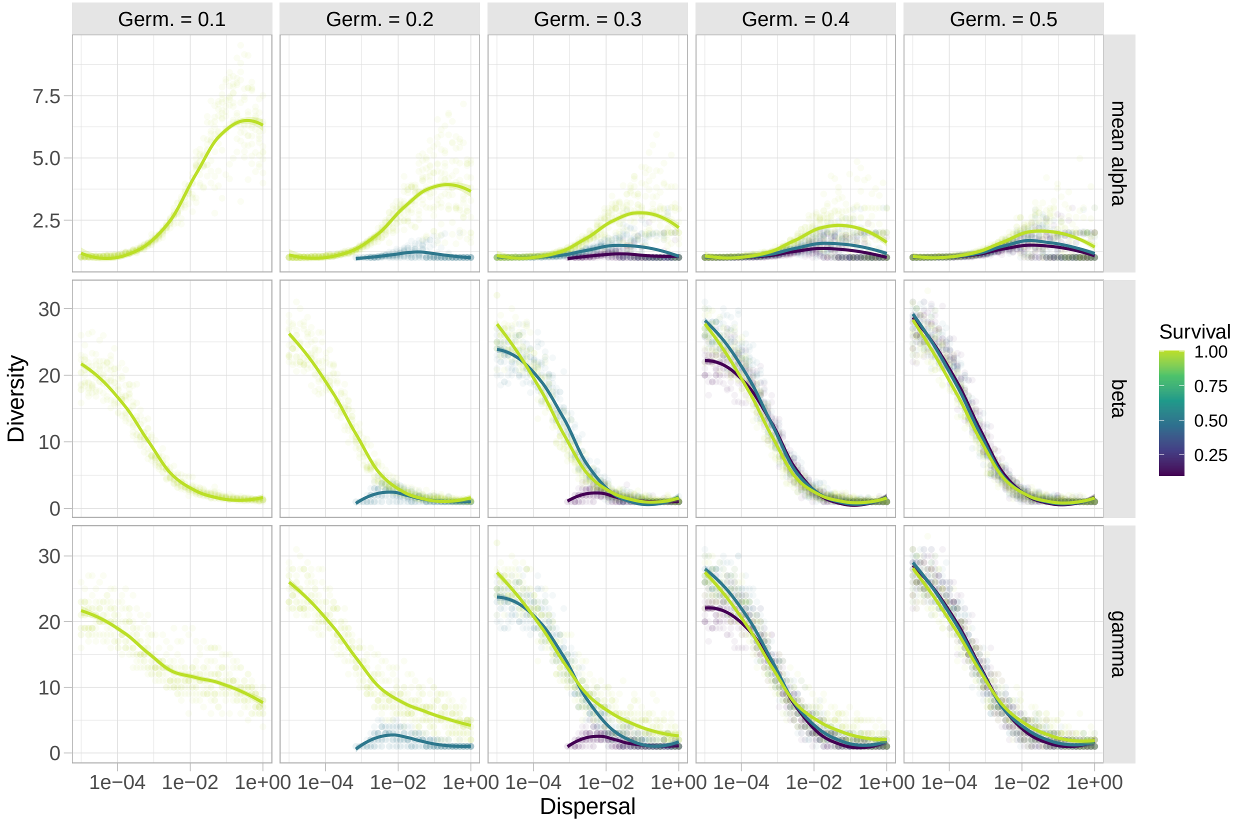


Figure S1: Dispersal-diversity relationships across a range of germination and survival rates with equal inter- and intraspecific competition coefficients. In these scenarios, lower survival rates eliminate combinations of low dispersal or germination that lead to extinctions. Higher survival allows diversity to persist across a range of dispersal and germination rates. The dispersal-dependent effects of survival are most noticeable when germination is low.

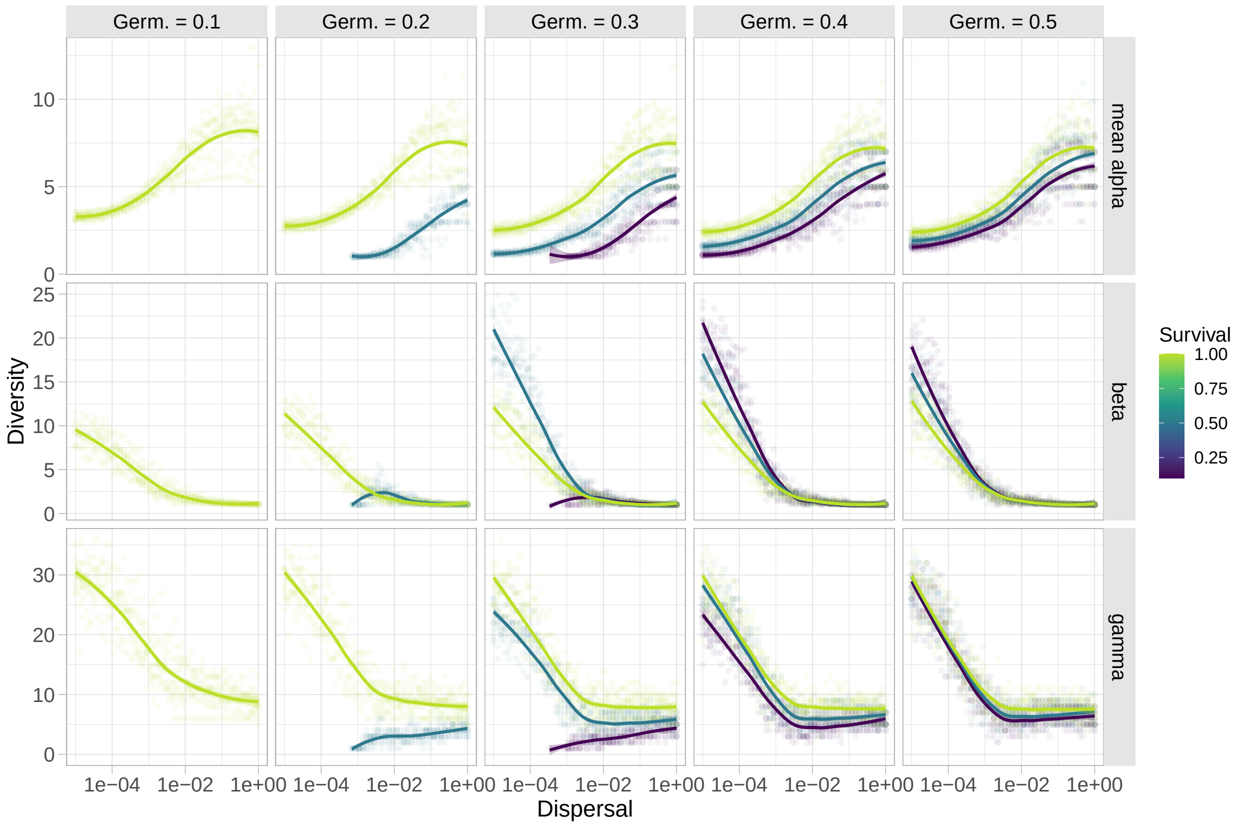


Figure S2: Dispersal-diversity relationships across a range of germination and survival rates with stable competition coefficients (i.e., intra > inter). In these scenarios, reduced survival has negative effects on alpha-, beta-, and gamma-diversity. But at low germination rates, survival must not be too low, otherwise no species persist in the metacommunity. Once germination is high enough (~ 0.5), the effect of survival on diversity is lower.
